# Waving goodbye to contrast: Self-generated hand movements attenuate visual sensitivity

**DOI:** 10.1101/474783

**Authors:** Madis Vasser, Laurène Vuillaume, Axel Cleeremans, Jaan Aru

## Abstract

It is well known that the human brain continuously predicts the sensory consequences of its own body movements, which typically results in sensory attenuation. Yet, the extent and exact mechanisms underlying sensory attenuation are still debated. To explore this issue, we asked participants to decide which of two visual stimuli was of higher contrast in a virtual reality situation where one of the stimuli could appear behind the participants’ invisible moving hand or not. Over two experiments, we measured the effects of such “virtual occlusion” on first-order sensitivity and on metacognitive monitoring. Our findings show that self-generated hand movements reduced the apparent contrast of the stimulus. This result can be explained by the active inference theory. Moreover, sensory attenuation seemed to affect only first-order sensitivity and not (second-order) metacognitive judgments of confidence.

## Introduction

Imagine having a drink in a pub. Your eyes fixate on the glass and your arm stretches to grab it. You are not bothered at all by the appearance of this long elongated shape (your arm) in your visual field. But now consider the same situation where a similar long elongated shape moves towards your beverage — be it a snake, or the arm of a colleague. In this case, you would immediately notice this potentially threatening moving object. Hence, it seems that the movement of our own body parts is processed differently from that of objects of the external world, many of which may nevertheless exhibit similar properties. In particular, it seems that the movement of our own body parts is not so salient, i.e. it captures less attention than the movement of external objects. In the present work we ask in which sense the movements of our own body are processed differently than that of external objects.

The brain’s ability to predict and attenuate the various sensory consequences of the movements of its own body is well known. However, the exact mechanisms underlying sensory attenuation are still debated (Blakemore et al., 1998; Bays et al., 2006; Brown et al., 2013; Clark, 2015). The active inference, or prediction error minimization theory (Friston, 2010; Hohwy, 2013; Clark, 2015) posits that the brain is constantly predicting sensory input. Predictions are then continuously compared with sensory input in such a way that only prediction errors are propagated further, with the overall computational goal of minimizing prediction error in the long term. Accordingly, performing actions is an efficient way of minimizing prediction errors by changing the sensory data so as to fit the predictions (Friston, 2010; Hohwy, 2013; Clark, 2015). The active inference theory suggests that movements are elicited by predicting the sensory consequences of the movement (e.g. that the hand will be moving in the visual field). These predictions will drive the behavior so that the organism will perform the necessary movements leading to the predicted state. However, the predicted consequences (for example that the hand will be moving towards the glass) are not in agreement with the current sensory data (where the hand is still in a resting position). The active inference theory proposes that this mismatch between the predictions and sensory data is resolved through withdrawal of attention from the current sensory input, resulting in sensory attenuation (for a longer treatment of this proposal see Brown et al., 2013; Clark, 2015). Hence, according to active inference sensory attenuation is a necessary counterpart of movement. This proposed mechanism of sensory attenuation can explain several findings in the literature that are difficult to understand under the classic efference copy view (e.g. Bays et al., 2006; Voss et al., 2008; van Doorn et al., 2014, 2015; Laak et al, 2017).

Recent work with virtual reality (VR) devices has brought direct support for active inference by demonstrating that attention is withdrawn from the area of the visual field where one’s own hand is currently moving (Laak et al, 2017). Participants were slower to detect stimulus changes (both movement and colour) in the condition where the change occurred behind their precisely tracked hand in VR. Importantly, participants themselves did not see their virtual hand, as it was rendered invisible. Thus, the effect was one of prediction rather than one of visual occlusion. Motor predictions also attenuate brain activity caused by corresponding visual feedback (Limanowski, Sarasso, & Blankenburg, 2018).

In the work by Laak et al (2017), which was designed to investigate active inference, the effect of lowered attention was observed through longer reaction times — an indirect measure of the quality of perception. In the current paper, we sought to extend these previous results to probe the effect of self-generated movement prediction on perception more directly. We based our study on the experimental approach used by Carrasco and colleagues (Carrasco, Ling & Read, 2004). In their paradigm, two Gabor patches with different orientations are presented, one cued and the other not, and the participants are asked to report the orientation of the Gabor patch with the stronger contrast. Through this experimental approach the authors could demonstrate that covert attention enhances the perceived contrast of an attended stimulus (Carrasco, Ling & Read, 2004). We reasoned that if 1) attention is withdrawn from the part of the visual field where one’s hand is moving (Laak et al., 2017) and 2) the deployment of attention affects perceived contrast (Carrasco et al., 2004), then the subjective contrast of objects in the region of the visual field where the hand is moving should be reduced.

In addition to first order sensitivity, we also sought to investigate the effect of self-generated movements on higher-order processes such as metacognition, that is, the ability to monitor and control one’s own mental states (Koriat, 2006). To come back to the pub setting, in addition to grabbing your beer, it is quite important to be sure that it really is yours, and that you are not actually stealing your neighbor’s beer, for instance. Metacognition is a crucial part of our daily life and a critical aspect of decision making in different domains (Metcalfe & Shimamura, 1996; Fleming, Dolan & Frith, 2012). In recent years, metacognition has become a prominent topic of investigation. However, to our knowledge, no previous study has directly addressed the influence of sensory attenuation on metacognition, and studies focusing on the interplay between attention and metacognition have so far yielded mixed results (Kanai, Walsh & Tseng, 2010; Sherman et al. 2015; Wilimzig & Fahle, 2008; Rahnev et al., 2011). Here, we measured the quality of metacognitive monitoring by asking participants to rate their confidence in their response on each trial, and by assessing the relationship between objective performance and subjective confidence (Galvin et al., 2003). This allowed us to explore our second research question: Do the self-generated movements also alter metacognitive accuracy?

To address these questions, we conducted two VR experiments where participants performed a visual two-alternative forced choice task coupled with moving their hand to overlap one of the target stimuli (Figure 1). Crucially, while the hand was tracked via sensors, the hand avatar was not shown to the participants. This made it possible to assess the extent to which visual expectations coupled to self-generated movement result in sensory attenuation.

**Figure 1.**
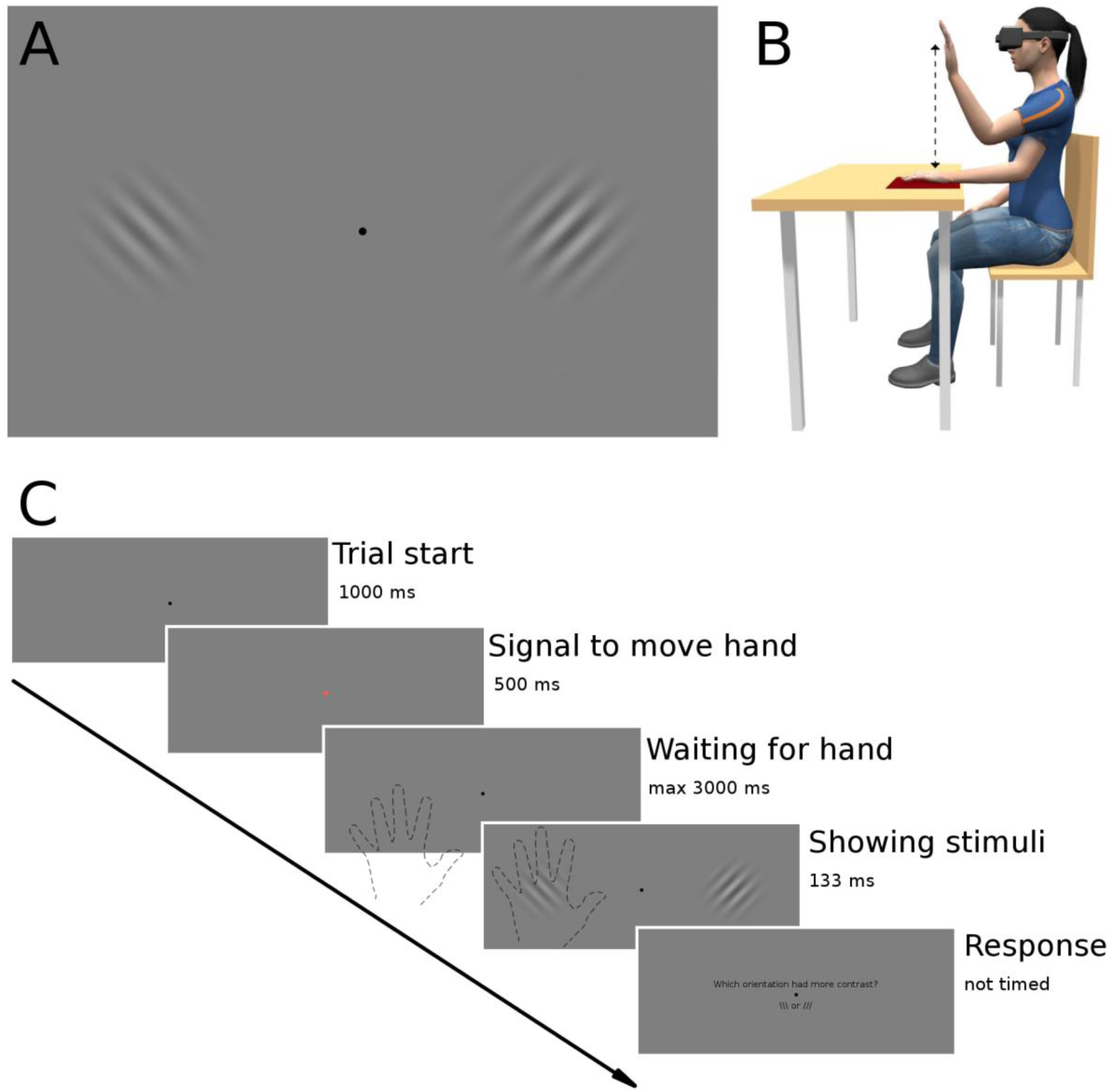
(A) Approximate example of the Gabor patches used as stimuli in the study. The pairs varied in contrast, frequency and orientation. The black dot in the middle is the gaze fixation point. (B) Physical setup of the experiment, with both rest and raised hand positions and approximate movement trajectory visualized. Reproduced with permission from Laak et al (2017). (C) General design for a single trial. Participants were instructed to perform a trained hand movement when the fixation cross changed color. This movement triggered the appearance of the two stimuli that were shown for 133 ms. The task consisted in reporting the orientation of the Gabor patch with the higher contrast. In Experiment 2, participants were additionally requested to report how confident they were in their decision on a scale from 1 (very unsure) to 4 (very sure). Note that participants’ hand was completely invisible to them. Hand outlines and Gabor patch sizes on the figure are illustrative.

In Experiment 1, we probed the first-order effects of hand movements on contrast judgements. Experiment 2 aimed to replicate Experiment 1 with a new group of participants, and to additionally probe the extent to which self-generated movements also influence metacognitive accuracy. In Experiment 2, participants thus also reported the confidence in their decisions, on every trial. A control condition in which the visual targets never appeared at locations situated behind the moving hand was also carried out. This allowed us to clarify whether the attenuation effect also extends to metacognitive ability or not.

## Methods

### Experimental setup

Our experimental setup was very similar to that of Laak et al (2017). The hardware solution consisted of an Oculus Rift CV1 virtual reality headset combined with a Leap Motion hand tracking device mounted directly on the goggles. This allowed us to have absolute control over the visual environment the participant perceived in the lab and also made it possible to record the relative transforms and parameters of the participant’s hands (position, velocity, orientation). Crucially, however, the hand avatar was not rendered in the virtual world. Thus, wherever we mention the "target behind the hand", the hand in question was completely invisible to the participant wearing the headset. As in Carrasco et al (2004), the visual stimuli consisted of Gabor patches with different contrasts, frequencies and orientations, shown in horizontally placed pairs (Fig. 1A). For the control condition of Experiment 2, the pairs were positioned vertically to avoid the effects of hand movement on contrast judgements.

Importantly, the participants’ task was orthogonal to the location of the stimuli (and thus to their hand movement) as their task was to report the orientation of the patch that seemed higher in contrast (see Carrasco et al., 2004) rather than to report its location (Experiment 1). In experiment 2, participants were additionally requested to report how confident they were in their decision. Stimulus presentation was triggered by the participants’ pre-trained hand movement. The stimuli were shown behind the hand only when the hand was moving upwards and the centre of the palm hit a virtual pre-defined target rectangle measuring 6 degrees in width and 3 degrees in height. Responses were registered by a left or right mouse click to indicate the orientation of the more contrasted Gabor and with the mouse scroll wheel to select a confidence rating from 1 to 4 followed by a mouse click to confirm the selected rating. We switched the hands used for movement and responses in the middle of the experiment so as to balance for possible preferences for one hand over the other. The stimulus marked as behind the hand was always on the side of the hand movement. A correctly executed hand movement only covered one of the Gabors. Participants were instructed to keep their gaze on the fixation point (a black dot in the middle of the field of view). As in Carrasco et al (2004) one of the two Gabor patches was always displayed with a set contrast (Standard), while the other was either of lower, identical, or higher in contrast (Target). In both experiments, we compared the percentage of trials reported to be higher in contrast in cases where the target appeared behind the invisible hand vs. cases where the target was not behind the hand. The temporal and spatial distribution of the targets in different conditions were kept constant in both experiments in relation to the participants’ hand movements.

### Apparatus

Participants were seated behind a table, with a standard computer mouse under one hand and an empty mouse pad under the other so as to promote consistent hand movements (Figure 1B). Participants were fitted with an Oculus Rift CV1 (Oculus VR, LLC; Oculus) VR headset with a refresh rate of 90 Hz, and a field of view of approximately 100 degrees and precise, low-latency positional tracking. The Leap Motion Controller (Leap Motion, Inc; Leap) Orion SDK (version 3.1.2) mounted on the headset was used to track hand movements. The Leap Motion sensor tracks the position, velocity, and orientation of hands and fingers with low latency and an average position accuracy of 1.2 mm (Weichert et al., 2013), with a field of view of 150 degrees horizontally and 120 degrees vertically. The experiments were conducted in a quiet and dark room.

### Stimuli

The default virtual environment consisted of a uniform gray background with a small black fixation dot (0.7 degrees in diameter) in the middle of the field of view, directly in front of the participant. The Gabor patches that appeared during hand movements were approximately 4 degrees in diameter from the perspective of the subject. Patches were generated only when the hand moved over a predefined rectangular target area in the virtual environment. The distance of the patches from the central fixation point ranged between 3-9 degrees to allow for some hand movement error in the horizontal axis. As in Carrasco et al. (2004), we varied the cycles per degree (CPD) of the gratings to avoid adaption to the stimuli and used 2 CPD and 4 CPD stimuli. The different contrasts and presentation times were chosen according to an extensive pilot study. The final contrasts corresponded to the following Michelson contrast values: 0.2; 0,24; 0.3; 0.36; 0.45. (Michelson contrast = (Lmax − Lmin) / (Lmax + Lmin), where Lmin is the lowest luminance of the stimulus and Lmax the maximum luminance of the stimulus.) The standard value used in all the trials was 0.3. Pairs of patches were shown for 133ms on each trial, after which participants reported the orientation of the patch that seemed to have more contrast. In Experiment 2, participants were additionally asked to report their confidence in their answer on a scale from 1 (very unsure) to 4 (very sure). Participants did not receive any feedback about their responses during the experiment.

### Participants

Participants were recruited through university lists and social media. A total of 54 healthy participants with normal or corrected-to-normal vision took part in the two experiments: 8 in the first (2 females, mean age 25), and 46 in the second experiment (27 females, mean age 23). In the second Experiment, 30 participants were assigned to the experimental group and 16 participants to the control group. All of the participants read and signed informed consent forms and participated in the experiments voluntarily. The VR experiments conducted in this study were approved by the Ethics Committee of the University of Tartu (Estonia) and the Ethics Committee of the Psychology Department of the Université Libre de Bruxelles (Belgium).

### Procedure

#### Instructions to the participants

The experiment was introduced to the participants as a study on attention. The hand tracking device was described as "a device that simply detects the general direction of hand movements that is needed to start each trial" in order to cover the true aim of the experiment. After signing an informed consent form, participants completed training on the required hand movement and on discriminating the Gabors. The exact protocol for hand movement training is explained elsewhere (Laak et al, 2017). Participants were informed that the Gabor patches would be triggered by the upwards movements of the hand.

#### Experiment 1

Each participant performed a total of 800 trials, of which 40% showed the Gabor with the stronger contrast behind the hand, 40% showed it at a location not behind the hand and 20% had equal contrast (standard vs standard). The conditions were balanced and randomized for each participant, and each participant was exposed to all of the conditions. After every 100 trials, a short rest of 10s occurred. The general design of a single trial for all of the experiments is shown in Figure 1C.

When participants missed the target area for the correct hand movement on several consecutive trials, they were verbally guided by the experimenter to improve their hand movement. After the experiment, participants answered control questions about complying with the instructions and were debriefed about the background and purposes of the study.

#### Experiment 2

In order to investigate the effects of sensory attenuation on metacognition, participants were randomly assigned to either an experimental group or a control group. Each participant performed a total of 400 trials in the experimental group and 300 trials in the control group. The experimental group was similar to Experiment 1. In the control group however, the pair of Gabor patches was shown vertically around the fixation dot instead of horizontally so as to make it so that neither stimulus appeared in the visual area where the hand was moving. In both groups participants were additionally asked to give their confidence in their decision on a scale from 1 (very unsure) to 4 (very sure).

### Data preprocessing

Data preprocessing and analysis were performed in R (version 3.1.2; R Core Team, 2015) using the afex package (Singmann et al., 2015), BayesFactor package using the medium default prior (Morey & Rouder, 2015) and ggplot2 package (Wikham, 2009). We discarded trials where the hand movement was not within the allowed constraints. Taken together, 56% of trials were rejected from Experiment 1, and 34% of trials from Experiment 2. This was due to the extreme precision necessary to ensure that hand movements were exactly in the same region of the visual field as the intended stimulus. Next we completely excluded participants that failed to show an increase in percentage of higher contrast judgements between the two lower test contrast values and the two higher test contrast values (hinting that the participant may have answered randomly). After this we were left with the following number of participants: 6 (Experiment 1) and 44 (Experiment 2).

### Statistical analysis

For statistical testing we used within-subject repeated measures analysis of variance (ANOVA) to test for the mean difference in the percentage of trials reported to be of higher contrast in different conditions. Degrees of freedom were corrected using the Greenhouse-Geisser method. Welch’s t-tests were used to test for the statistical difference between accuracy, confidence and metacognitive ability in the experimental group compared to the control group in Experiment 2 (the equal contrast trials were not included in this analysis since accuracy in this case cannot be computed). Metacognitive ability was estimated through the type-II area under the receiver-operating curve (AROC), which determines the rate of correct and incorrect responses at each confidence level (Kornbrot, 2006). A second approach for evaluating metacognitive ability was fitting a mixed logistic regression model of accuracy with group, confidence and their interaction as fixed effects, and a random participant intercept. The regression slope was taken as an indicator of metacognitive ability (see Siedlecka, Paulewicz & Wierzchoń, 2016). We also used Bayesian statistics to assess the likelihood that data was in favour of the alternative or null hypothesis using the default medium prior of the BayesFactor R package (Morey & Rouder, 2015). This is especially relevant to interpret non-significant p-values in conventional statistics (Dienes, 2014). Bayes Factors (BFs) above 1 indicate evidence for the alternative hypothesis whereas BF below 1 indicate evidence for the null hypothesis.

## Results

### Experiment 1

Using a motion tracking device in combination with VR technology we were able to investigate whether self-generated movements influence visual perception despite participants never seeing their hand (Figure 1). Participants performed a two-alternative forced choice task where they had to report the orientation of the Gabor patch with the higher contrast. Crucially for the experiments, one of the Gabor patches was located in the same visual area as the self-generated hand movement.

The main results are illustrated in Figure 2, which suggests that for all contrast differences, target items that appear behind the hand are less often reported to be of higher contrast than targets that appear at the other location, which is indicative of sensory attenuation. A within-subjects ANOVA showed a main effect of contrast (F(1.37, 6.85) = 48.12, p < 0.001, η_p_^2^ = 0.80) and importantly, also a strong effect of whether the stimulus appeared behind the invisible hand or not (F(1,5) = 15.07, p = 0.01, η_p_^2^ = 0.58). We found no interaction between condition and contrast (F(1.79,8.97) = 3.1, p = 0.10, η_p_^2^ = 0.07). This suggests that apparent contrast is indeed reduced when shown in the area of the visual field where the hand is currently moving.

**Figure 2.**
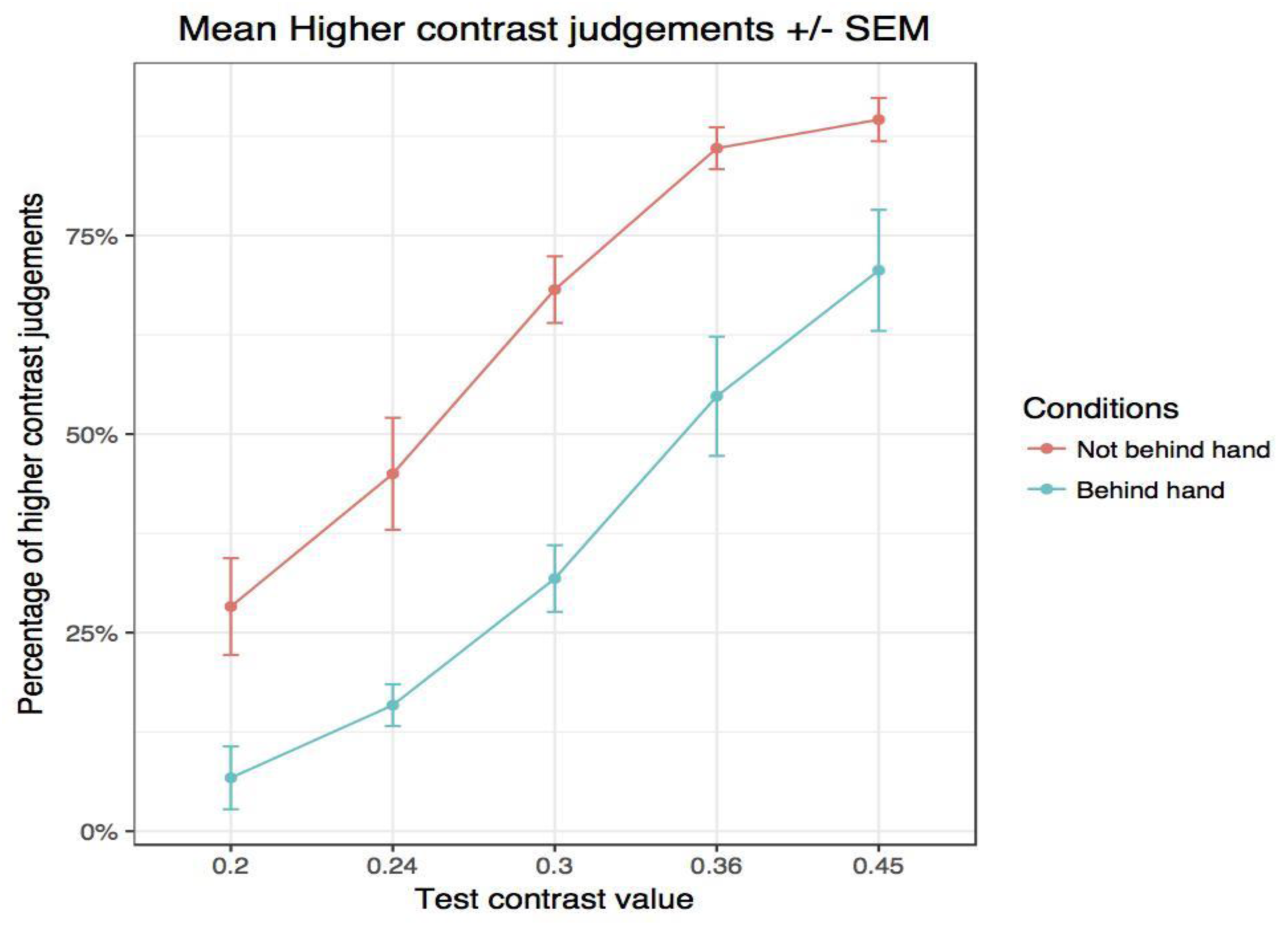
Results of Experiment 1. X axis is the value of the test contrast, Y axis denotes the percentage of trials that were reported as higher in contrast. Blue line shows the target stimulus appearing behind the hand and red line shows the target stimulus not behind the hand.

### Experiment 2

The first goal of Experiment 2 was to replicate Experiment 1 in another laboratory. Additionally, Experiment 2 was also aimed at exploring the extent to which sensory attenuation influences metacognitive processes. The design of Experiment 2 was thus identical to that of Experiment 1, except that participants were also requested to report the confidence in their judgement on every trial, using a scale ranging from 1 (no confidence) to 4 (full confidence). Experiment 2 included a control condition in which targets never appeared behind the moving hand.

#### Experimental group

##### Replication of experiment 1

We first sought to replicate our findings from Experiment 1. The same analysis confirmed these results, by revealing a main effect of contrast (F(1.57,45.39) = 212.10, p < 10^−4^, η_p_^2^ = 0.66) and a significant difference in the percentage of “higher contrast” responses between the "behind the hand" condition and the "not behind the hand" condition (F(1, 29) = 11.13, p = 0.002, η_p_^2^ = 0.18). There was again no interaction between Condition and Contrast (F(3.60,104.27) = 1.01, p = 0.40, η_p_^2^ = 0.005). Similarly to Experiment 1, this indicates that apparent contrast is reduced when shown in the same visual area as the self-generated movement.

##### Confidence

An analysis of variance revealed, as expected, a main effect of contrast on confidence (F(3.24,93.99) = 55.49, p < 10^−4^, η_p_^2^ = 0.17), as well as a significant difference in confidence ratings between the "behind the hand" condition and the "not behind the hand" condition (F(1,29) = 6.17, p = 0.02, η_p_^2^ = 0.004). The interaction between Contrast and Condition was also significant (F(2.05,59.42) = 10.12, p = 0.0001, η_p_^2^ = 0.05) (Figure 3). This suggests that participants were able to adjust their confidence according to the perceived contrasts.

**Figure 3.**
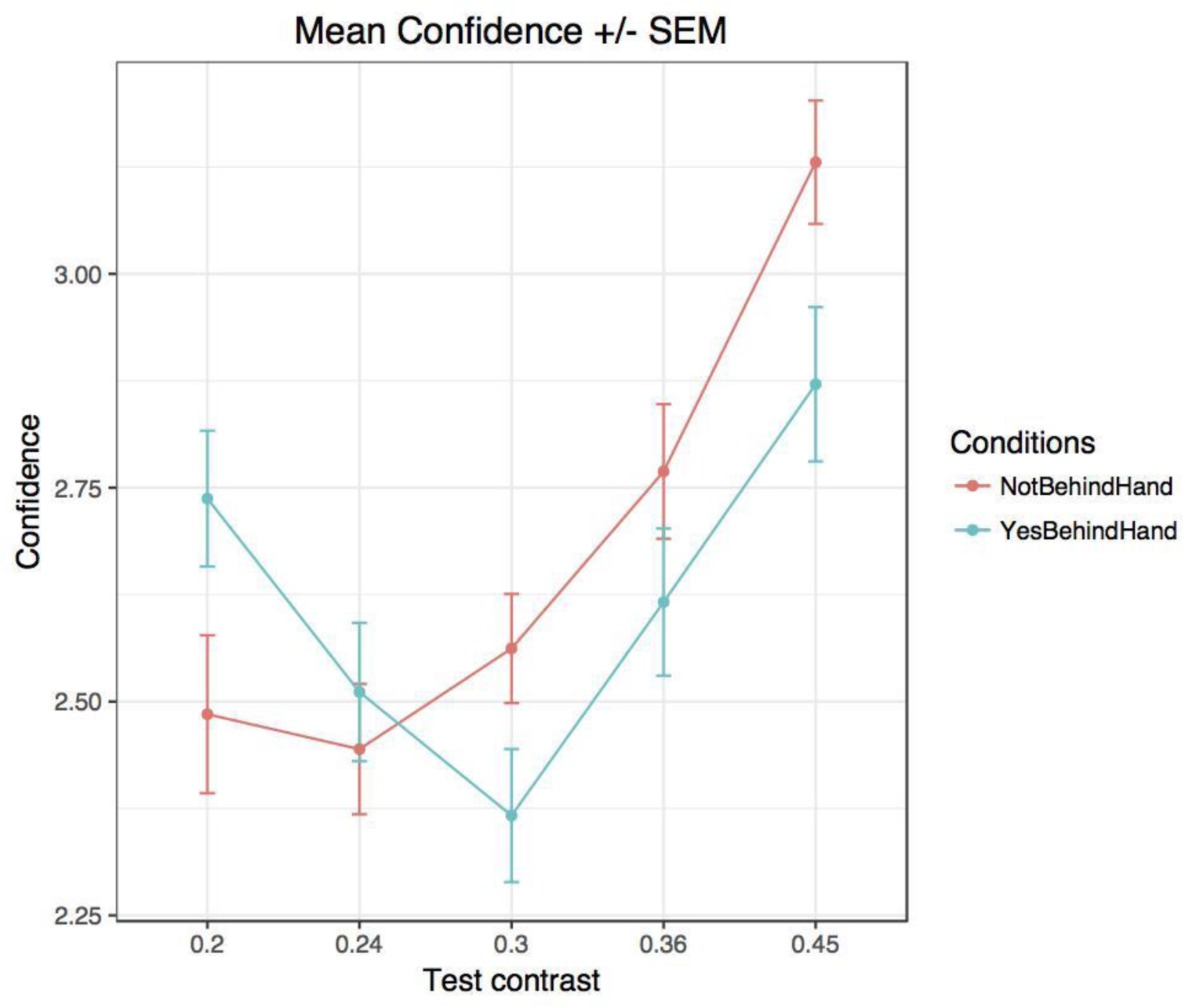
Mean confidence ratings in Experiment 2 (experimental group) as a function of condition (Blue curve: Target stimulus behind the hand; Red curve: Target stimulus not behind the hand). Percentages of trials that were reported as seen to be higher in contrast between the two conditions were similar to Experiment 1.

In order to evaluate the effect of self-generated movements on metacognitive ability we added a control group. Indeed, in the experimental group one of the stimuli was always behind the hand movement (the test contrast or the standard contrast) and thus there was always one of the two contrasts that appeared to be reduced. To investigate the effect of sensory attenuation on metacognitive ability it was necessary to compare the experimental group to a control group in which both of the stimuli were shown in a different visual area than the self-generated hand movement. To do so, in the control group the stimuli were presented vertically around the fixation point instead of horizontally. This made it possible to use exactly the same procedure as used in the experimental group but with none of the stimuli presented in the visual area behind which the hand was moving.

#### Control group

We first controlled that participants in the control group performed the task correctly. Analysis of variance also revealed a main effect of contrast on the percentage of stimuli reported to be higher in contrast (F(1.45,18.87) = 133.71, p < 10^−4^, η_p_^2^ = 0.90) and on confidence ratings (F(3.01,39.17) = 19.24, p < 10^−4^, η_p_^2^ = 0.27).

#### Comparison between the experimental group and the control group

We found no effect of group on type 1 sensitivity (d’: t(26.95) = -0.36, p = 0.72) or criterion (t(34.56) = 0.93, p = 0.36) and no effect of group on confidence ratings either (t(30.57) = 0.21, p = 0.84).

##### Metacognition

Metacognitive accuracy as measured with the AROC did not differ between groups (t(21.54) = -0.49, p = 0.63, BF = 0.35)(Figure 4) nor did Type II bias (BROC: t(35.40) = 0.68, p = 0.50, BF = 0.36). The mixed logistic regressions between accuracy and confidence, with the regression slope taken as an indicator of metacognitive ability (see Siedlecka, Paulewicz & Wierzchoń, 2016), yielded similar results (no difference between the control group and the experimental group slopes: estimate = -0.02, *z* = −0.38, *p* = 0.70).

**Figure 4.**
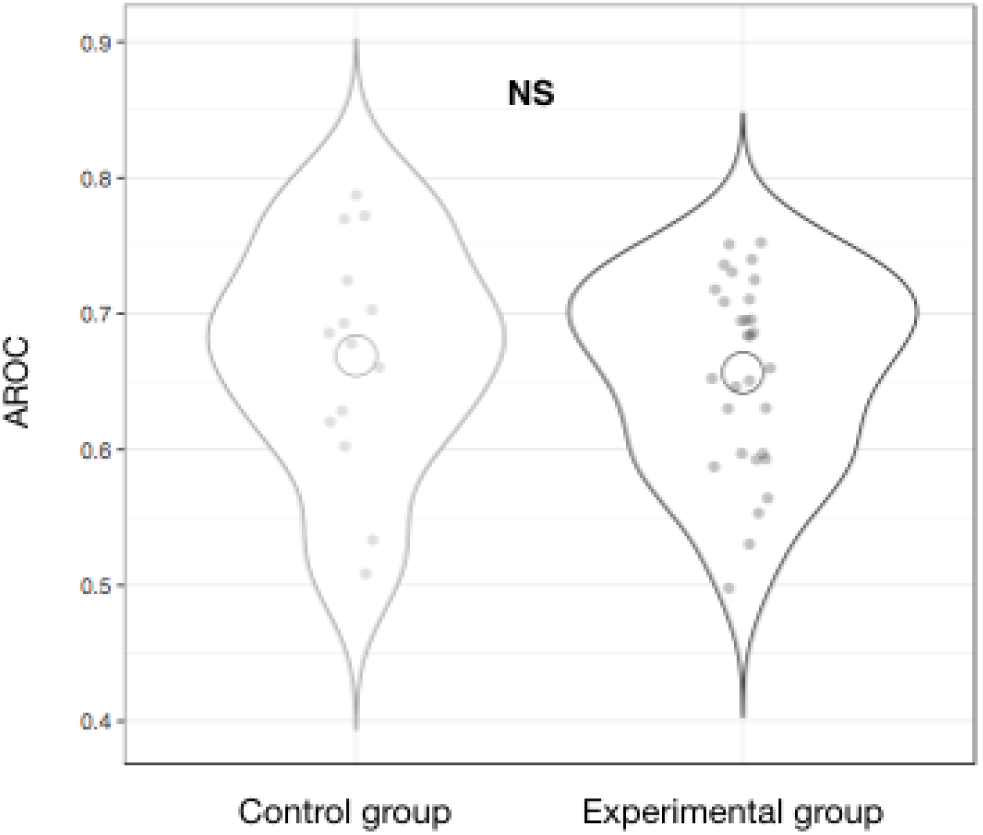
The mean AROC in Experiment 2 for the control group and the experimental group.

## Discussion

In two independent experiments, we observed that (invisible) self-generated movements influence visual sensitivity while leaving metacognitive accuracy intact. This suggest that perception is directly influenced by one’s expectations about the sensory consequences of one’s own movements. In our paradigm visual targets that would normally be occluded by our own moving limbs were perceptually attenuated although these targets were fully visible.

This was achieved by combining VR goggles and a hand tracking device, allowing us to present stimuli in the same visual area as the hand and at the same time being able to keep the hand invisible to the participants (Laak et al., 2017). This novel paradigm allowed us to investigate sensory attenuation in the visual domain, which until recently was often overlooked due to the technical limitations associated with the challenge of removing visual sensory feedback.

After initiating the hand movement, participants had to indicate the orientation of the Gabor patch with higher contrast. We observed that the contrast of the stimulus behind the invisible hand was significantly reduced. We replicated these results in an independent group of participants and estimated whether metacognitive ability would also be affected by our experimental manipulation. In order to measure metacognitive accuracy, participants were additionally required to report their confidence in their response on each trial. There was no difference in metacognitive accuracy between the experimental group and the control group, suggesting that participants in the experimental group were able to adjust their subjective confidence to their objective performance. Indeed, even though the contrast of one of the stimuli was reduced through sensory attenuation, they could equally well judge their confidence in their decision. That is, if a low test contrast was attenuated they were more confident in their decision - they could tell that it was easier - and, similarly, if a high test contrast was attenuated they were less confident in their decision. This is in line with previous work from Kanai, Walsh & Tseng (2010) who showed that metacognitive ability was preserved when attention was diverted using several methods as well as the study from Sherman et al. (2015) who found no difference in metacognitive ability between a full attention and a diverted attention condition in a perceptual task. Furthermore, being able to tell that there is sensory attenuation might be a crucial cue in recognizing an action as being internally triggered and not externally triggered. This has been emphasized in the study of schizophrenia patients who showed deficit in attenuating the sensory consequences of their actions which in turn gave rise to issues in discriminating self-generated actions from externally triggered events (Blakemore et al., 2003; Shergill et al., 2005; Fletcher & Frith, 2009). Sensory attenuation thus appears to only influence first-order processes.

The main results confirm those of Laak et al. (2017) but go a step further by indicating that self-generated movements do not only impact reaction times through attenuating attention, they also influence visual sensitivity per se. Our results thus offer support for the active inference account, which posits sensory attenuation of visual processing due to sensory precision in being reduced during movement (Friston, 2010; Hohwy, 2013; Clark, 2015).

### Alternative explanations to the present findings

Active inference can readily explain our present results: in order to move, precision of the sensory data that would be in conflict with the predicted outcome has to be reduced and this is done by withdrawing attention from the parts of the visual field where the hand is currently moving. The consequence of withdrawn attention is that the Gabor that would be on the path of the moving hand (if the hand would be visible) is seen with less contrast. According to the active inference theory, sensory attenuation is the consequence of withdrawing attention from the sensory data coming from the *specific parts* of the visual field where the hand is moving.

However, in principle one could also explain the present findings with a more general account of attentional suppression in space. In particular, one could claim that when a limb is moving, visual spatial attention is more generally withdrawn, i.e. not only from the trajectory where the hand is moving. From our present results we know that it cannot be a suppression of the whole visual field (otherwise we could not obtain differences between the two conditions), but for example it could be claimed that in the hemifield where the hand is currently moving attention is withdrawn. We think that there are several reasons to doubt such a general suppression. First, although we did not test for the spatial specificity in this study, in the previous work with the similar setup (Laak et al., 2017; experiment 2) we observed that attention seems not to be withdrawn from locations that are in the same visual hemifield as the hand movement but not directly on the movement path. Second, such a general suppression would be disadvantageous: For example, if one is reaching for an object this general attentional suppression would lead to the withdrawal of attention from the reach target. This would be unreasonable and in fact experimental data has demonstrated that attention is enhanced around reach targets (Rolfs et al., 2013). The last finding might prompt the question whether it is in conflict with the present results: How can reach targets be attentionally prioritized while the hand movement trajectory is attenuated? Active inference theory can explain this as enhancing the precision of the target is a separate process which can occur independently of and together with the reduction of precision from those parts of the visual field where the hand is moving.

The more classic efference copy theory gives a more specific alternative explanation to the present results than the general suppression account. Efference copy theory suggests that sensory input that is predicted by the own motor command is cancelled. Such forward models are learned over the lifetime and can be used to explain sensory attenuation (e.g. Blakemore et al., 1998). However, this theory has also trouble explaining the present results. In particular, subjects cannot have been learning over the lifetime that the hand movement predicts the appearance of a Gabor. In principle, one could suggest that the association between hand movement and appearance of Gabors could have been learned over the course of the experiment, but this cannot explain the difference between the two conditions (as all hand movements were followed by the presentation of *both* Gabors). More generally, the efference copy theory cannot account for several findings on the field that are readily explained by active inference theory (for overview see Brown et al., 2013; Clark, 2015). Hence, overall, active inference casts the most comprehensive explanation of sensory attenuation both for the current findings and for the variety of results available in the literature.

### Limitations and further questions

Due to the limits of the used hardware, it was impossible to record participant’s eye movements during the experiments. Therefore, no technical guarantee can be given that the participants maintained stable gaze on the fixation point throughout the experiment. To mitigate this, we had targets with varying contrast on both sides of the visual field, so participants had no incentive to prefer one side over the other. The participants were also instructed to keep their eyes on the fixation point during the trials and all verbally complied. Future research methodology could benefit from the next generation VR headsets and add-on devices rapidly becoming available at a reasonable price that allow eye-tracking (e.g. FOVE, 2018; Tobii, 2018).

Prospective investigations are also required to examine more precisely how our findings generalise over different categories of stimuli or stimulus feature differentiation. It has been shown that these aspects can interact with attention and metacognition (Stein & Peelen, 2017; Matthews et al., 2018). In addition, the precise influence of evidence-reliability remains to be elucidated in more extensive work as it has also been shown to affect confidence judgments and metacognition (Boldt, De Gardelle & Yeung, 2017; Bang & Fleming, 2018; Denison & al., 2018).

Using this paradigm, one could further investigate the characteristics of visual attenuation caused by self-generated hand movements. Indeed, it would be interesting to compare our results with a static version, in which participants just hold their hand at the target position all the time. According to the active inference theory, we would expect that without any movement the attenuation would disappear. One could also investigate the role of agency and intention to act by removing the self-generated component and having the experimenter move the hand of the participant (Limanowski et al., 2018). Another way to investigate this would also be to test whether subjects who are less prone to sensory attenuation also exhibit diminished agency judgements. Furthermore, we would expect attenuation to build up with stronger expectations of hand movement over time (see Voss et al., 2008 and Bays et al., 2006). The role of proprioception could be tested as well using a vibrator on the arm tendon in order to induce proprioceptive noise and disrupt these signals. Similarly, inducing a rubber hand illusion (Botvinick & Cohen, 1998) has also been shown to lead to cases of sensory attenuation (Burin et al., 2017; Burin et al., 2018). Using this paradigm in that context could make it possible to better understand the role of body ownership and to assess the extent to which the illusion could misplace the effect of the self-generated movement on another visual area. Finally, the perceptual consequences of own hand movements have been established over decades of learning and cannot be "unlearned" with only a few hundred trials. It would be interesting to test how sensory attenuation develops for example for a newly learned tool.

## Conclusions

We conducted two experiments showing that self-generated hand movements reduce the subjective contrast of objects in the part of the visual field where the hand is directly moving, but do not alter metacognitive monitoring. The present experimental paradigm provides a novel way to study sensory attenuation and demonstrates the usefulness of modern VR tools for investigating fundamental questions about the computations that run our lives.

## Acknowledgements

We thank Tõnis Kristian Koppel for invaluable help with programming and the reviewers for improving our manuscript.

## Funding

This work was supported by PUT1476 and IUT20-40 of the Estonian Research Council and an European Research Council Advanced Grant RADICAL to A.C. A.C. is a Research Director with the F.R.S.-FNRS (Belgium). JA was also supported by the European Union’s Horizon 2020 Research and Innovation Programme under the Marie Skłodowska-Curie grant agreement no. 799411.

### Data availability

All data and codes are available from the authors upon request.

## References

Bang, D., & Fleming, S. M. (2018). Distinct encoding of decision confidence in human medial prefrontal cortex. Proceedings of the National Academy of Sciences, 115(23), 6082–6087.

Bays, P. M., Flanagan, J. R., & Wolpert, D. M. (2006). Attenuation of self-generated tactile sensations is predictive, not postdictive. PLoS biology, 4(2), e28.

Blakemore, S. J., Wolpert, D. M., & Frith, C. D. (1998). Central cancellation of self-produced tickle sensation. Nature neuroscience, 1(7), 635.

Blakemore, S. J., Oakley, D. A., & Frith, C. D. (2003). Delusions of alien control in the normal brain. Neuropsychologia, 41(8), 1058–1067.

Boldt, A., De Gardelle, V., & Yeung, N. (2017). The impact of evidence reliability on sensitivity and bias in decision confidence. Journal of experimental psychology: human perception and performance, 43(8), 1520.

Botvinick, M., & Cohen, J. (1998). Rubber hands ‘feel’ touch that eyes see. Nature, 391(6669), 756.

Brown, H., Adams, R. A., Parees, I., Edwards, M., & Friston, K. (2013). Active inference, sensory attenuation and illusions. Cognitive processing, 14(4), 411–427.

Burin, D., Pyasik, M., Salatino, A., & Pia, L. (2017). That’s my hand! Therefore, that’s my willed action: How body ownership acts upon conscious awareness of willed actions. Cognition, 166, 164–173.

Burin, D., Pyasik, M., Ronga, I., Cavallo, M., Salatino, A., & Pia, L. (2018). “As long as that is my hand, that willed action is mine”: Timing of agency triggered by body ownership. Consciousness and cognition, 58, 186–192.

Cardoso-Leite, P., Mamassian, P., Schütz-Bosbach, S., & Waszak, F. (2010). A new look at sensory attenuation: Action-effect anticipation affects sensitivity, not response bias. Psychological science, 21(12), 1740–1745.

Carrasco, M., Ling, S., & Read, S. (2004). Attention alters appearance. Nature neuroscience, 7(3), 308.

Clark, A. (2015). Surfing uncertainty: Prediction, action, and the embodied mind. Oxford University Press.

Denison, R. N., Adler, W. T., Carrasco, M., & Ma, W. J. (2018). Humans incorporate attention-dependent uncertainty into perceptual decisions and confidence. Proceedings of the National Academy of Sciences, 115(43), 11090–11095.

Dienes, Z. (2014). Using Bayes to get the most out of non-significant results. Frontiers in psychology, 5, 781.

Fleming, S. M., Dolan, R. J., & Frith, C. D. (2012). Metacognition: computation, biology and function.

Fletcher, P. C., & Frith, C. D. (2009). Perceiving is believing: a Bayesian approach to explaining the positive symptoms of schizophrenia. Nature Reviews Neuroscience, 10(1), 48.

Friston, K. (2010). The free-energy principle: a unified brain theory?. Nature Reviews Neuroscience, 11(2), 127.

FOVE, Inc. (2018). Eye Tracking VR dev kit. Retrieved June 17, 2018, from https://www.getfove.com.

Galvin, S. J., Podd, J. V., Drga, V., & Whitmore, J. (2003). Type 2 tasks in the theory of signal detectability: Discrimination between correct and incorrect decisions. Psychonomic Bulletin & Review, 10(4), 843–876.

Hohwy, J. (2013). The predictive mind. Oxford University Press.

Kanai, R., Walsh, V., & Tseng, C. H. (2010). Subjective discriminability of invisibility: a framework for distinguishing perceptual and attentional failures of awareness. Consciousness and cognition, 19(4), 1045–1057.

Koriat, A. (2006). Metacognition and consciousness. Institute of Information Processing and Decision Making, University of Haifa.

Kornbrot, D. E. (2006). Signal detection theory, the approach of choice: Model-based and distribution-free measures and evaluation. Perception & Psychophysics, 68(3), 393–414.

Laak, K. J., Vasser, M., Uibopuu, O. J., & Aru, J. (2017). Attention is withdrawn from the area of the visual field where the own hand is currently moving. Neuroscience of Consciousness, 3(1).

Limanowski, J., Sarasso, P. and Blankenburg, F. (2018), Different responses of the right superior temporal sulcus to visual movement feedback during self- generated vs. externally generated hand movements. European Journal of Neuroscience, 47(4), 314–320.

Matthews, J., Schröder, P., Kaunitz, L., van Boxtel, J. J., & Tsuchiya, N. (2018). Conscious access in the near absence of attention: critical extensions on the dual-task paradigm. Phil. Trans. R. Soc. B, 373(1755), 20170352.

Metcalfe J, Shimamura AP. (1996) Metacognition. Cambridge, MA: MIT Press.

Morey, R. D., Rouder, J. N., Jamil, T., & Morey, M. R. D. (2015). Package ‘BayesFactor’. URL< http://cran.r-project.org/web/packages/BayesFactor/BayesFactor.pdf> (accessed10.06. 15).

R Core Team (2015). A language and environment for statistical computing. R Foundation for Statistical Computing, Vienna, Austria. URL https://www.R-project.org/.

Rahnev, D., Maniscalco, B., Graves, T., Huang, E., De Lange, F. P., & Lau, H. (2011). Attention induces conservative subjective biases in visual perception. Nature neuroscience, 14(12), 1513.

Rolfs, M., Lawrence, B. M., & Carrasco, M. (2013). Reach preparation enhances visual performance and appearance. Phil. Trans. R. Soc. B, 368(1628), 20130057.

Shergill, S. S., Samson, G., Bays, P. M., Frith, C. D., & Wolpert, D. M. (2005). Evidence for sensory prediction deficits in schizophrenia. American Journal of Psychiatry, 162(12), 2384–2386.

Sherman, M. T., Seth, A. K., Barrett, A. B., & Kanai, R. (2015). Prior expectations facilitate metacognition for perceptual decision. Consciousness and Cognition, 35, 53–65.

Schwarz, K.A., Pfister, R., Kluge, M., Weller, L., & Kunde, W. (2018). Do we see it or not? Sensory attenuation in the visual domain. Journal of Experimental Psychology: General, 147(3), 418.

Siedlecka, M., Paulewicz, B., & Wierzchoń, M. (2016). But I was so sure! Metacognitive judgments are less accurate given prospectively than retrospectively. Frontiers in psychology, 7, 218.

Singmann, H., Bolker, B., Westfall, J., Højsgaard, S., Fox, J., & Lawrence, M. (2015). afex: Analysis of factorial experiments. R package version 0.13–145.

Stein, T., & Peelen, M. V. (2017). Object detection in natural scenes: Independent effects of spatial and category-based attention. Attention, Perception, & Psychophysics, 79(3), 738–752.

Tobii, AB. (2018). Tobii Pro VR Integration. Retrieved June 17, 2018, from https://www.tobiipro.com/product-listing/vr-integration/.

Van Doorn, G., Hohwy, J., & Symmons, M. (2014). Can you tickle yourself if you swap bodies with someone else? Consciousness and cognition, 23, 1–11.

Van Doorn, G., Paton, B., Howell, J., & Hohwy, J. (2015). Attenuated self-tickle sensation even under trajectory perturbation. Consciousness and cognition, 36, 147–153.

Voss, M., Ingram, J. N., Wolpert, D. M., & Haggard, P. (2008). Mere expectation to move causes attenuation of sensory signals. PLoS One, 3(8), e2866.

Weichert, F., Bachmann, D., Rudak, B., & Fisseler, D. (2013). Analysis of the accuracy and robustness of the leap motion controller. Sensors, 13(5), 6380–6393.

Wickham, H. (2009). ggplot2: elegant graphics for data analysis. Springer New York, 1(2), 3.

Wilimzig, C., Tsuchiya, N., Fahle, M., Einhäuser, W., & Koch, C. (2008). Spatial attention increases performance but not subjective confidence in a discrimination task. Journal of Vision, 8(5), 7–7.

